# Modeling predicts differences in CAR T cell signaling due to biological variability

**DOI:** 10.1101/2022.01.14.476364

**Authors:** Vardges Tserunyan, Stacey D. Finley

## Abstract

In recent decades, chimeric antigen receptors (CARs) have been successfully used to generate engineered T cells capable of recognizing and eliminating cancer cells. The structure of CARs frequently includes costimulatory domains, which enhance the T cell response upon antigen encounter. However, it is not fully known how the CAR co-stimulatory domains influence T cell activation in the presence of biological variability. In this work, we used mathematical modeling to elucidate how the inclusion of one such co-stimulatory molecule, CD28, impacts the response of a population of engineered T cells under different sources of variability. Particularly, our simulations demonstrate that CD28-bearing CARs mediate a faster and more consistent population response under both target antigen variability and kinetic rate variability. We identify kinetic parameters that have the most impact on mediating cell activation. Finally, based on our findings, we propose that enhancing the catalytic activity of lymphocyte-specific protein tyrosine kinase (LCK) can result in drastically reduced and more consistent response times among heterogeneous CAR T cell populations.

## 1 INTRODUCTION

T cells engineered to express a chimeric antigen receptor (CAR) have emerged as a novel tool for combatting cancer by generating an immune response against cancer cells. The key step in this immunotherapeutic approach is to produce T cells expressing an artificially designed CAR, which activates the T cell upon encountering a target cancer cell (Met et al., 2019). To achieve this goal, CARs feature an antigen recognition domain derived from a single-chain variable fragment (scFv) of a monoclonal antibody specific to the antigen of interest, while their cytoplasmic portion includes different combinations of intracellular signaling domains (Akhoundi et al., 2021). The first generation of CARs has a single cytoplasmic CD3ζ signaling domain attached to the recognition domain by a transmembrane linker. Later advances in CAR design resulted in second-generation CARs, which have one additional costimulatory signaling domain originating from signaling endodomains of natively occurring costimulatory receptors (Srivastava and Riddell, 2015). For example, the CD28 costimulatory domain is utilized in two of the FDA-approved CAR-T therapies as of February 2021 (Albinger et al., 2021). Notably, the inclusion of CD28 significantly enhances proliferation and target cytotoxicity among the T cells (Zhao et al., 2015). However, despite this progress, certain gaps in CAR-T performance remain. Importantly, current CAR-T therapies are sufficiently effective only against liquid tumors, while the performance against solid tumors is still limited (Sterner and Sterner, 2021). In addition, CD28-based CAR-T therapies suffer from some major side effects (Namuduri and Brentjens, 2016), which can be life-threatening (Tabernero and Thompson, 2018). These considerations necessitate further improvements in the CAR-T technology, with two broad aims: on one hand, make CAR-T cells efficient against a more extensive set of targets; on the other hand, mitigate the observed side effects.

Mathematical modeling is an integral approach in Systems Biology and has been applied to explain biological phenomena and generate testable hypotheses to guide experimental research (Kitano, 2002). One successful strategy of mathematical modeling is ordinary differential equation (ODE)-based mechanistic modeling, whereby dynamic processes occurring in a biological system are represented via differential equations describing the rate of change of each component as a function of its interactions with other components (Raue et al., 2013). With suitable estimates of interaction parameters and initial conditions, these ODEs can be integrated to obtain time courses for each component. In the past, various research groups, including ours, have utilized ODE-based mechanistic modeling to gain quantitative insights into immune cell signaling dynamics. For example, this approach was successful in describing native T cell receptor (TCR)-induced activation of MAPK/ERK signaling (Altan-Bonnet and Germain, 2005). Our group has focused on simulating the dynamics of ERK signaling in CAR-T cells in response to antigen binding (Rohrs et al., 2020). Particularly, by calibrating model parameters against data obtained by phosphoproteomic measurements, the authors were able to obtain a close correspondence between model predictions and observed experimental dynamics. In addition, we made quantitative comparisons between first-generation CAR-T cells with CD3ζ as the only signaling domain and second-generation CAR-T cells with an additional CD28 co-stimulatory domain. Thus, modeling was successfully used to gain quantitative insights into CAR-T cells, thereby augmenting experimental knowledge.

Variability has been observed for essentially all dimensions of single cell measurements, with the ensemble behavior of a cell population not necessarily reflecting the behavior of an individual cell (Altschuler and Wu, 2010). Examples of variability caused by differential protein expression have been observed among endogenous T cell populations, including T regulatory cells (Yuan et al., 2014), T helper cells, and T killer cells alike (Zhang et al., 2019). Thus, it is natural to expect that various sources of variability would affect the performance of CAR-T cells used in a therapeutic setting. Although we previously used our calibrated model of CAR-mediated signaling to investigate the effects of variability in the expression levels of signaling proteins on ERK activation (Cess and Finley, 2020), other manifestations of variability remain to be addressed. First, while in prior single-cell simulations we assumed that the cell encounters a well-defined antigen concentration on the surface of the target, this is not true in the therapeutic setting. For example, prior measurements of acute lymphoblastic leukemia (ALL) B cells found a range of values for the cell surface concentration of CD19 (Nerreter et al., 2019), the signature antigen targeted by the recognition domain of many CAR-T cells. Second, the chemical reactions that mediate signal transduction are subject to fluctuations not only due to stochastic protein expression, but also variability in cell state, local microenvironment and number of molecular collisions (McAdams and Arkin, 1999; Harton and Batchelor, 2017). Thus, in our current work, we set out to investigate and compare the performance of first-generation and CD28-bearing second-generation CAR-T cells under two modes of heterogeneity: exposure to stochastic concentrations of the target antigen and stochastic kinetic rates of signaling processes. Then, we used a data-driven approach to quantify the importance of different kinetic parameters for determining the ERK activation time of cells. Finally, based on these findings, we propose strategies to further enhance the efficiency of CAR-T cells in a therapeutic setting.

## 2 METHODS

### CAR-induced ERK signaling model

The ODE-based model utilized in our work was developed in MATLAB by our research group (Rohrs et. al, 2020). The model includes four signaling modules: phosphorylation of ITAM regions of the CD3ζ domain in response to antigen binding, inhibitory activity of CD45 and SHP1, LAT signalosome formation and MAPK signaling (**Supp. Fig. S1**). The model was calibrated on experimental data and gives accurate quantitative estimates of the levels of signaling species, indicating that it constitutes a plausible description of the underlying biological processes. Particularly, the model gives a mechanistic explanation for the increased cytoplasmic concentration of doubly-phosphorylated ERK (ppERK) in response to antigen binding to the CAR. We use ppERK as the primary model output, as it mediates cell activation. Given the non-transient “all-or-nothing” response typically displayed by ppERK in response to antigen stimulation (Altan-Bonnet and Germain, 2005), we termed all cells with more than half of their total ERK pool in doubly phosphorylated form as “active”, with the time needed to reach this state termed the “activation time”.

### Monte Carlo simulations

In simulations of variable antigen concentration encountered by CAR T cells, we assumed the antigen distribution (in units of molecules/µm^2^) to be lognormal with the scale parameter µ=1.0 and the scatter parameter σ=0.5. Our choice of the lognormal distribution was based on the fact that most intracellular protein abundances closely obey a lognormal distribution (Furusawa, et al., 2005). We picked the location and scale parameters to match the observed concentration of CD19, a frequent target of CAR T cell therapies. The surface concentration of CD19 ranges between 0.16 molecules/µm^2^ and 5.2 molecules/µm^2^ in ALL B cells (Nerreter, et al., 2019). Simulations for variability of kinetic parameters were carried out by sampling each parameter from an independent normal distribution with the mean equal to the parameter’s accepted value and a standard deviation equal to a third of the mean. These simulations were repeated for different antigen concentrations: 4.5 molecules/µm^2^ (“low”, close to experimentally observed) and 45 molecules/µm^2^ (“high”, near saturating). All Monte-Carlo simulations were run in MATLAB, with multiple random seeds to ensure reproducibility.

### Gradient-boosted tree predictor

The gradient-boosted tree ensemble is a non-linear machine learning method used successfully for both regression and classification tasks. It is based on a succession of individually weak decision trees. The first tree in the sequence fits the observed outcome directly, while each successive tree fits the residual left from the collective prediction of its predecessors. Thus, together, these decision trees achieve a significantly enhanced performance over a single decision tree (James et al., 2013). We utilized a *scikit-learn* implementation of gradient-boosted trees (Pedregosa et al., 2011) to obtain a data-driven mechanism-independent predictor of cell activation time based on the values of kinetic parameters as sole input. The training and testing dataset was generated by using the ODE-based model to compute ERK activation times for 10^5^ different sets of values of 48 impactful kinetic parameters. Then, the predictor was trained on this synthetic dataset in *Python 3*.*7* to regress cell activation time from the corresponding parameter vector. The accuracy of the gradient-boosted tree ensemble was adjusted by tuning hyperparameters and evaluating resultant performance by 5-fold cross validation of the coefficient of determination (R^2^) and explained variance (EV). Notably, by analyzing the formulas for these metrics, it can be shown that if the obtained R^2^ and EV are equal, then the predictor is unbiased.

### Permutation importance scores

The permutation feature importance score is defined to be the decrease in the R^2^ value of model prediction when a single feature value is randomly shuffled across all data points (Pedregosa et al., 2011). This shuffling procedure preserves the marginal distribution of the feature across the population, but otherwise decouples it from the output value of the data point. The underlying assumption is that if a feature is important in determining the output value, then this population-wide random shuffling will greatly reduce the predictive power of the model resulting in a proportionate drop in predictor performance as evaluated by R^2^ values on a test set. Similarly, since this decoupling procedure is highly unlikely to improve the performance of a predictor, permutation importance scores cannot be negative. We utilized the *scikit-learn* library to calculate permutation importance scores with five different shufflings for each parameter, using these iterative calculations to evaluate the statistical significance of obtained scores according to a one-tailed *t-*test.

### Parameter selection by optimization

Having quantified the importance of kinetic parameters on ERK activation time, we had the goal of isolating a handful of kinetic parameters whose manipulation would result in the largest reduction of ERK activation time. For this purpose, we picked as candidates the five highest-scoring kinetic parameters among first-generation cells in low-antigen conditions, excluding the affinity constant between the CAR and the antigen (to focus on targeting catalytic activities). Next, we used particle swarm optimization to minimize the following objective function

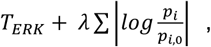

where *T*_*ERK*_ is ERK activation time, *p*_*i*_ is the parameter value to be optimized, *p*_*i,0*_ is the default value of that parameter, and λ is a user-defined constraint parameter that specifies how much the ratio of the optimal and default parameter value is weighted. We varied λ across a range. The structure of the objective function, inspired by the LASSO parameter selection technique, allows us to optimize parameters with the goal of decreasing activation time while changing as few parameters as possible. Particularly, it is long established that the penalization of the sum of absolute values in classic LASSO results in parameter selection by which only a subset of parameters is assigned non-zero values, while the rest remain at zero (Tibshirani, 1996). Since, in our case, the “default” value of the parameter is non-zero, the use of the logarithm of fold change allows us to assign zero penalty to unchanged values, while penalizing any change in proportion to the default value. We hypothesized that this manner of penalization would result in a similar parameter selection such that ERK activation times would be reduced by manipulating as few parameters as possible.

## 3 RESULTS

### 3.1 Population response with variable antigen exposure

We simulated the response of CAR-T cell populations to a distribution of antigen concentrations. Thus, 10^5^ cells expressing either the first-or second-generation CARs were stimulated by an antigen concentration coming from a lognormal distribution *in silico*, and their activation times were recorded. Particularly, we predicted the number of CAR-T cells that became active in the course of the simulation (**Fig. 1A**) and summarized their activation times in a histogram (**Fig. 1B**). The presence of the CD28 costimulatory domain in the second-generation CAR resulted in a higher percentage of activated cells (82.59% vs 99.97% for first- and second-generation CAR-T cells, respectively), with a shorter mean activation time (18.53 min vs. 8.93 min for first- and second-generation cells, respectively). Additionally, the distribution of the population response has a smaller standard deviation among second-generation cells compared to the first-generation cells (3.00 min vs 5.42 min, respectively). This indicates that the CD28 domain not only confers greater activation in a shorter time, but also provides for a more consistent response. These effects of the CD28 domain can be attributed to the shifted dose response curve (**Supp. Fig. S2**).

**Figure 1.**
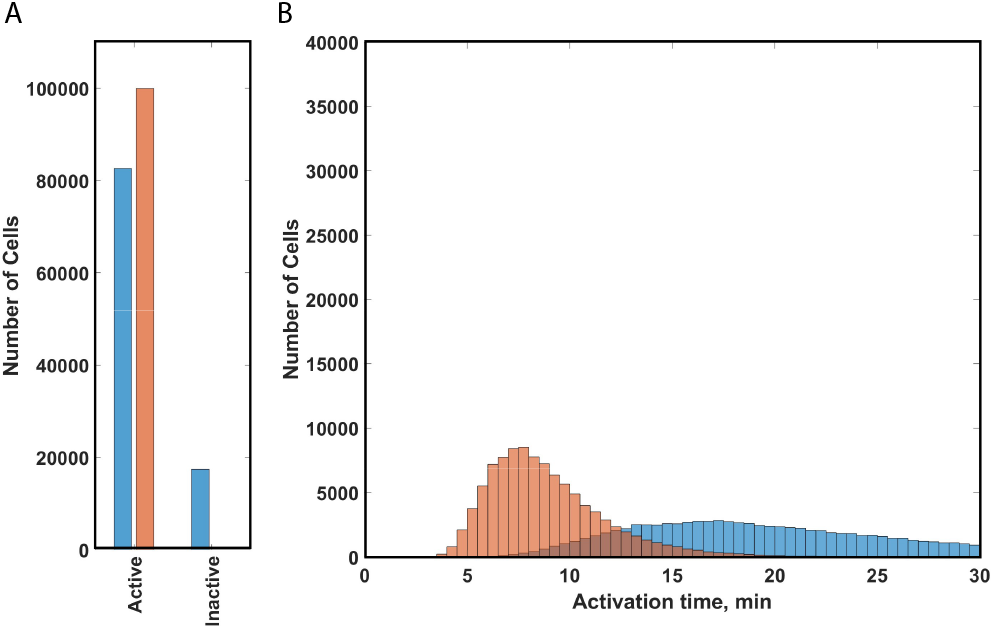
Activation of CAR T cells exposed to varying amounts of antigen. Predicted activation for 105 cells with first- and second-generation CAR constructs stimulated by the same stochastic antigen concentrations in silico: (A) The number of cells becoming active or remaining inactive in course of the 30-minute simulation (no inactive cells were detected among second-generation cells); (B) Distribution of the activation times for cells that became active in course of the 30-minute simulation. *Blue*, predictions for first-generation cells (CAR-CD3ζ); *orange*, predictions for second-generation cells (CAR-CD3ζ-CD28).

### 3.2 Population response with variable kinetic parameters

Another source of variability we set out to explore is the variability in effective rates of the reactions involved in signal transduction. Various sources of biological noise can result in heterogeneity of effective kinetic rates across a genetically uniform population. Thus, it is important to compare the performance of cells engineered with first- and second-generation CARs given variable kinetic parameters to account for the consequences of this mode of heterogeneity. To investigate the effects of kinetic variability, we performed simulations for first- and second-generation cells with randomized values of 48 influential parameters in conditions of either “low” or “high” antigen exposure. As evidenced by the resulting population distributions, the inclusion of the CD28 domain results in shorter mean activation time and a smaller standard deviation (**Fig. 2**). Particularly, in low antigen conditions, more cells were activated in cells expressing the CAR that contains the CD28 domain (76.68% vs. 94.24%, for first- and second-generation cells, respectively), with a shorter activation time (14.06 min vs. 8.91 min, for first- and second-generation cells, respectively) and reduced standard deviation (6.48 min vs. 5.15 min, for first- and second-generation cells, respectively). In high antigen conditions, the presence of CD28 caused a similar change in the population response, albeit less pronounced (98.41% vs. 98.91% of cells were activated, with mean activation time of 4.45 min vs 3.32, and standard deviation of 2.83 min vs 1.95 min, for first- and second-generation cells, respectively).

**Figure 2.**
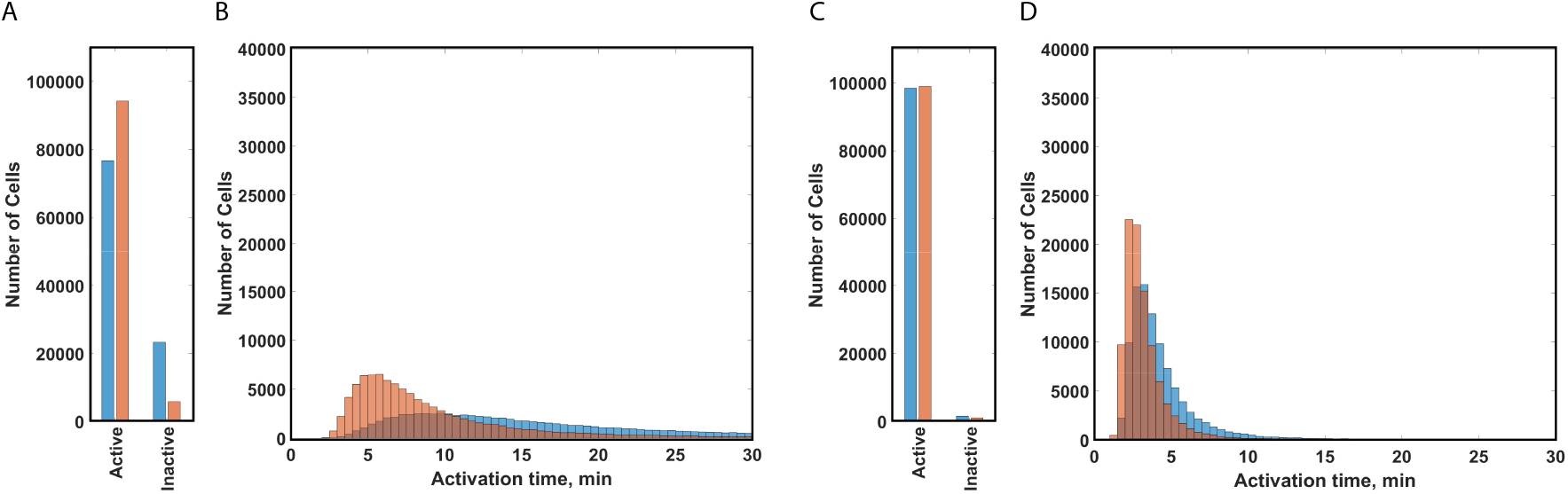
Activation of CAR T cells with varied kinetic parameters. Predicted activation for 105 cells with first-or second-generation CAR constructs stimulated by the same antigen concentration (*high* or *low*) with stochastic kinetic parameters *in silico*: (A) The number of cells becoming active or remaining inactive in course of the 30-minute simulation with low antigen stimulation; (B) Distribution of the activation times for cells from panel A that became active in course of the 30-minute simulation; (C) The number of cells becoming active or remaining inactive in course of the 30-minute simulation with high antigen stimulation; (D) Distribution of the activation times for cells from panel C which became active in course of the 30-minute simulation. *Blue*, predictions for first-generation cells (CAR-CD3ζ); *orange*, predictions for second-generation cells (CAR-CD3ζ-CD28).

### 3.3 Mechanism of CD28 induced changes in population behavior

A natural question to pursue is to determine the mechanism by which CD28 causes this reduction in the mean and standard deviation of response times. Specifically, we focused on three possible mechanisms: the interaction between CD28 and the adaptor protein Grb2, the interaction between CD28 and the adaptor protein GADS (Watanabe et al., 2006), and the enhancement of the catalytic activity of LCK by CD28 (Rohrs et al., 2018). The baseline model includes all three of these CD28 mechanisms (**Supp. Fig. S1**). To isolate the contribution of each interaction, we again ran simulations with stochastic kinetic parameters but with only one CD28-mediated mechanism available at a time. We found that in the case of the isolated CD28/Grb2 interaction, second-generation cells perform poorly compared to first-generation cells, with increased mean activation time and standard deviation (Fig. 3A; Table 1). Similar results were obtained for the case of the isolated CD28/GADS interaction (**Fig. 3B**; Table 1). On the other hand, with the isolated effect of CD28 on LCK catalytic activity, we saw the reduced mean and standard deviation that are the hallmark of second-generation cells (**Fig. 3C**; Table 1). Thus, based on our simulations, the kinetic effect of CD28 on LCK activity is the leading mechanism of CD28’s role in influencing ERK activation. Overall, the interaction between CD28 and Grb2 or the interaction between CD28 and GADS proved insufficient to bring about any improvement, while CD28’s effect on LCK’s catalytic activity is shown to be necessary and sufficient.

**Figure 3.**
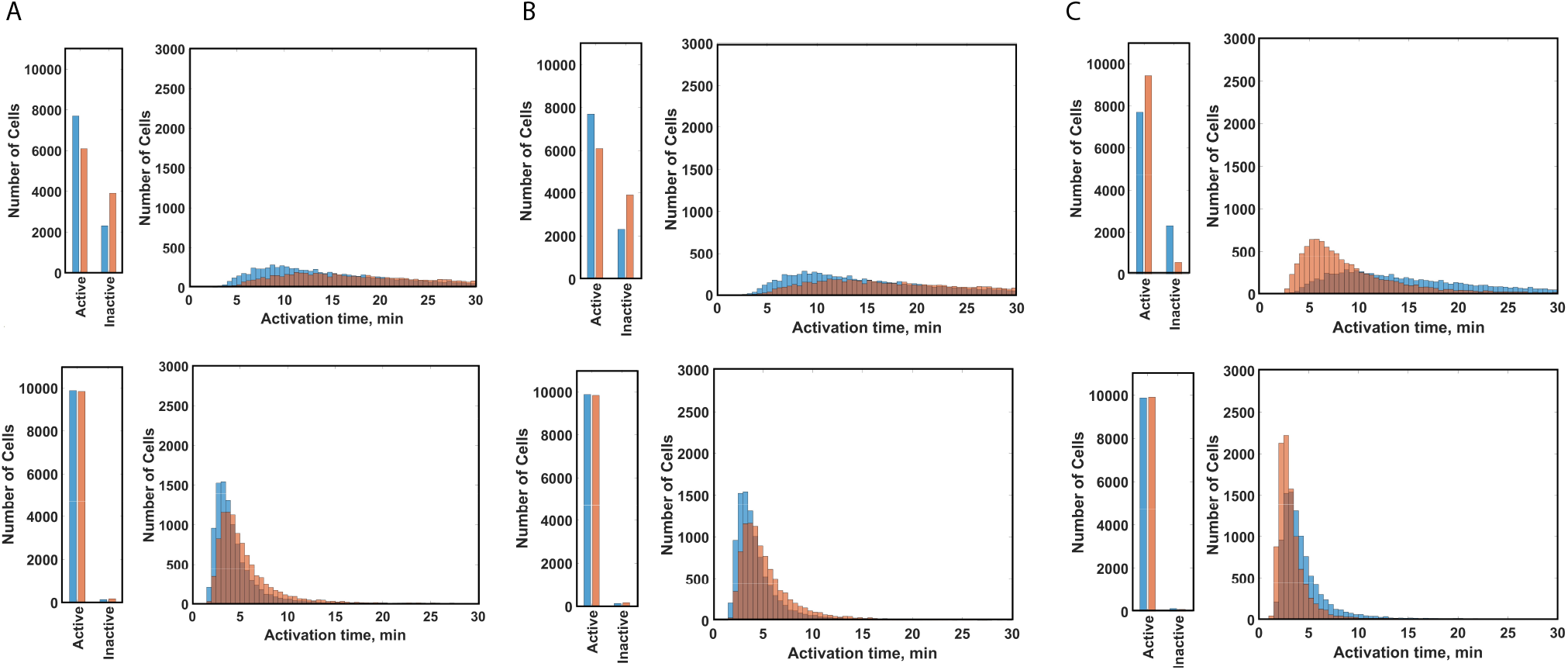
Activation of CAR T cells considering alternative CD28 signaling mechanisms. Predicted activation for 105 cells with first- and second-generation constructs with one CD28 signaling mechanism implemented at a time. (A) CD28 exclusively associates with Grb2; (B) CD28 exclusively associates with GADS; (C) CD28 exclusively affects the activity of LCK. Bar plot shows the number of cells becoming active or remaining inactive in course of the 30-minute simulation. Histogram shows the distribution of the activation times for cells that became active in the 30-minute simulation. *Top*, low-antigen simulation. *Bottom*, high-antigen simulation: *Blue*, predictions for first-generation cells (CAR-CD3ζ); *orange*, predictions for second-generation cells (CAR-CD3ζ-CD28).

### 3.4 Data-driven sensitivity analysis of activation time

Next, we aimed to quantify the impact of each parameter through a data-driven procedure. Here, the influence of each parameter is exclusively based on the model output without explicit consideration of the mechanistic details of the model. Such data-driven analysis would help identify potential targets for enhancing cell activation. In order to obtain data-driven importance scores for those 48 parameters, we first created a synthetic dataset in which the model was simulated for 10^5^ different and independently sampled values of the 48 parameters. This created a 48-by-10^5^ matrix of parameter values with corresponding activation times. This procedure was performed for four settings: first- and second-generation cells, each in “low” and “high” antigen conditions. Then, we developed a gradient-boosted tree ensemble (GBTE) and trained it on each dataset. Here, each of the 48 varied parameters were treated as input features and activation time as the output value. Hyperparameters were tuned until satisfactory predictive performance by 5-fold cross-validation was obtained. Two performance metrics of the resulting GBTE, R^2^ and EV, are given for each condition in Table 2. Since the obtained values for R^2^ and EV are identical in all settings, this implies that the GBTE is an unbiased estimator.

With a successful “black-box” predictive model at hand, we set out to quantify the importance of each parameter for the prediction made by the GBTE. We did this by using permutation importance scores. The importance of each parameter was quantified in each of the four conditions (**Fig. 4, Supp. Fig. S3**). With these results, we isolated the top five kinetic parameters that have a large impact on first-generation cells in low-antigen conditions: the catalytic activity of LCK in phosphorylating ITAM regions of CD3ζ (*Kcat_LCKPU_CD3z*), association rate of CSK with LCK (*CSKon*), catalytic activity of ZAP70 (*Kcat_ZAP*), catalytic activity of CD45 in dephosphorylating LCK (*Kcat_CD45_LCK505*), and the catalytic activity of CD45 in dephosphorylating ITAM regions of CD3ζ (*Kcat_CD45_A1*). Notably, the three highest-scoring parameters were identical between first- and second-generation cells in low antigen conditions.

**Figure 4.**
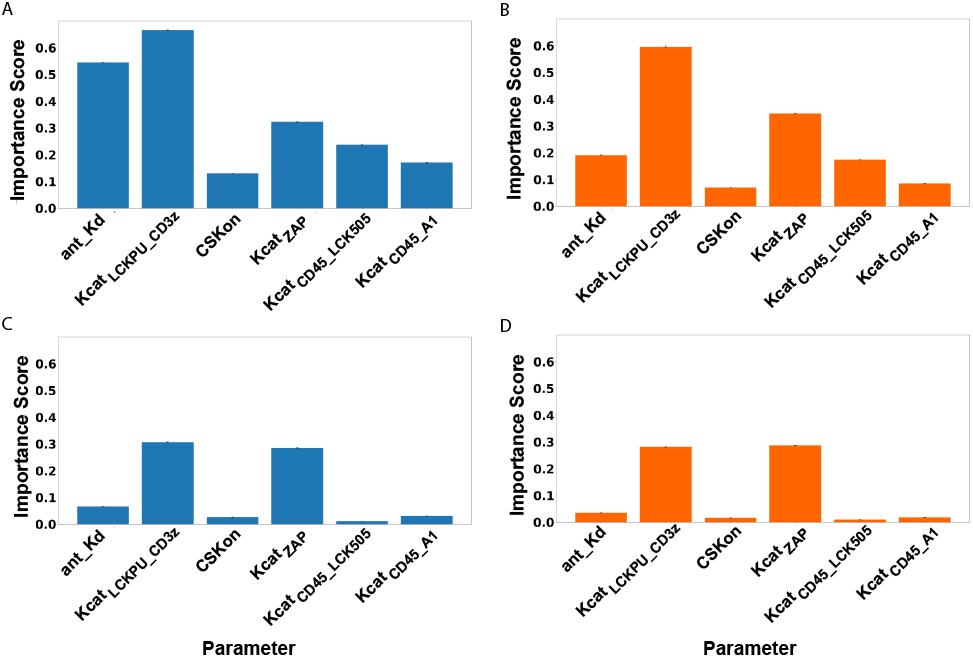
Permutation importance scores for select kinetic parameters used in machine learning model to predict cell activation times. A gradient boosted tree was used to predict the cell activation times based on model kinetic parameters. We show the most important kinetic parameters that influence the predicted activation time under different conditions: (A) CAR-CD3ζ with low antigen concentration, (B) CAR-CD3ζ-CD28 with low antigen concentration, (C) CAR-CD3ζ with high antigen concentration, (D) CAR-CD3ζ-CD28 with high antigen concentration.

### 3.5 Parameter selection by constrained optimization

We hypothesized that due to the large impact in determining the activation time of the cell, each of the five influential parameters identified from the GBTE could serve as a target for engineering more efficient CAR-T cell lineages. A key goal of such engineering would be to minimize the number of interventions into the system given the complexity of designing proteins with desired properties and then expressing them in engineered cells. Thus, we performed parameter selection by constrained optimization to identify which one of these parameters could serve as the most optimal target of a limited experimental intervention. The objective function was to minimize activation time while modifying as few of parameters as possible (see Methods section). This optimization routine was performed with both first- and second-generation cells, each in low- and high-antigen conditions, and with different values of the optimization constraint parameter (**Fig. 5**). When the value of the constraint parameter is 10, kinetic parameters do not change at all in course of the optimization. However, as we relax the constraint parameter (i.e., reduce its value to 1.0), two kinetic parameters change within one order of magnitude to actuate a decrease in cell activation time: *Kcat_LCKPU_CD3z* and *Kcat_ZAP*. These two kinetic parameters remain the only ones whose values should be optimized even when we further decrease the constraint parameter. Thus, we predict that when the goal is to decrease cell activation time, an increase in *Kcat_LCKPU_CD3z* and *Kcat_ZAP* will result in the largest such decrease.

**Figure 5.**
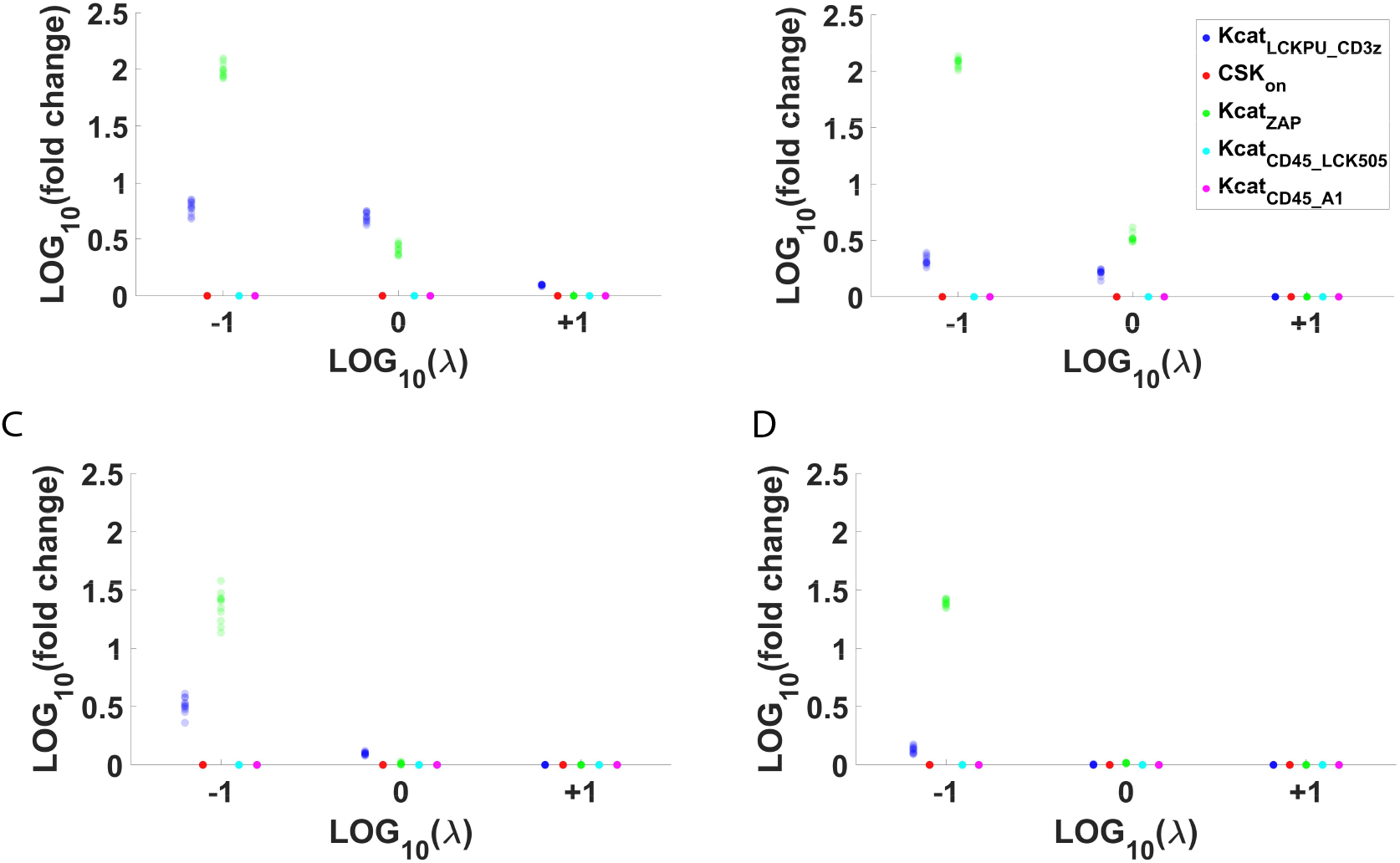
Results from constrained optimization. Logarithms of fold changes in each of the five parameters used in the constrained optimization to minimize activation time are shown for given constraint strength (*λ*), under different conditions: (A) first-generation cells under low antigen; (B) second-generation cells under low antigen; (C) first-generation cells under high antigen; (D) second-generation cells under high antigen.

To test the consequences of targeting the parameters identified by the optimization procedure, we again performed simulations with varying antigen concentrations, with one or both of the selected parameters set to their optimized values and all others kept at their default values. Based on our results, setting *Kcat_ZAP* to the optimized value resulted in the reduction of activation time (**Fig. 6A**), with a greater fraction of second-generation cells becoming active compared to first-generation cells (Table 3). However, setting *Kcat_LCKPU_CD3z* to the optimized value was sufficient to not only induce a drastic reduction in cell activation time, but also to make first-generation cells more efficient than second-generation cells (**Fig. 6B, 6C**; Table 3). Particularly, with *Kcat_LCKPU_CD3z* optimized alone, both populations showed 100% activation with mean activation times 5.20 min vs. 6.25 min for first- and second-generation cells, respectively; when both *Kcat_LCKPU_CD3z* and *Kcat_ZAP* were optimized, both populations showed 100% activation with mean activation times of 3.09 min vs. 3.54 min for first- and second-generation cells, respectively (Table 3). When we reran simulations of kinetic variability with the same optimized parameters, we obtained similar results. Specifically, *Kcat_LCKPU_CD3ζ* optimization is predicted to be sufficient to make first-generation cells respond faster to antigen presence than second-generation cells (**Fig. 7**; Table 4).

**Figure 6.**
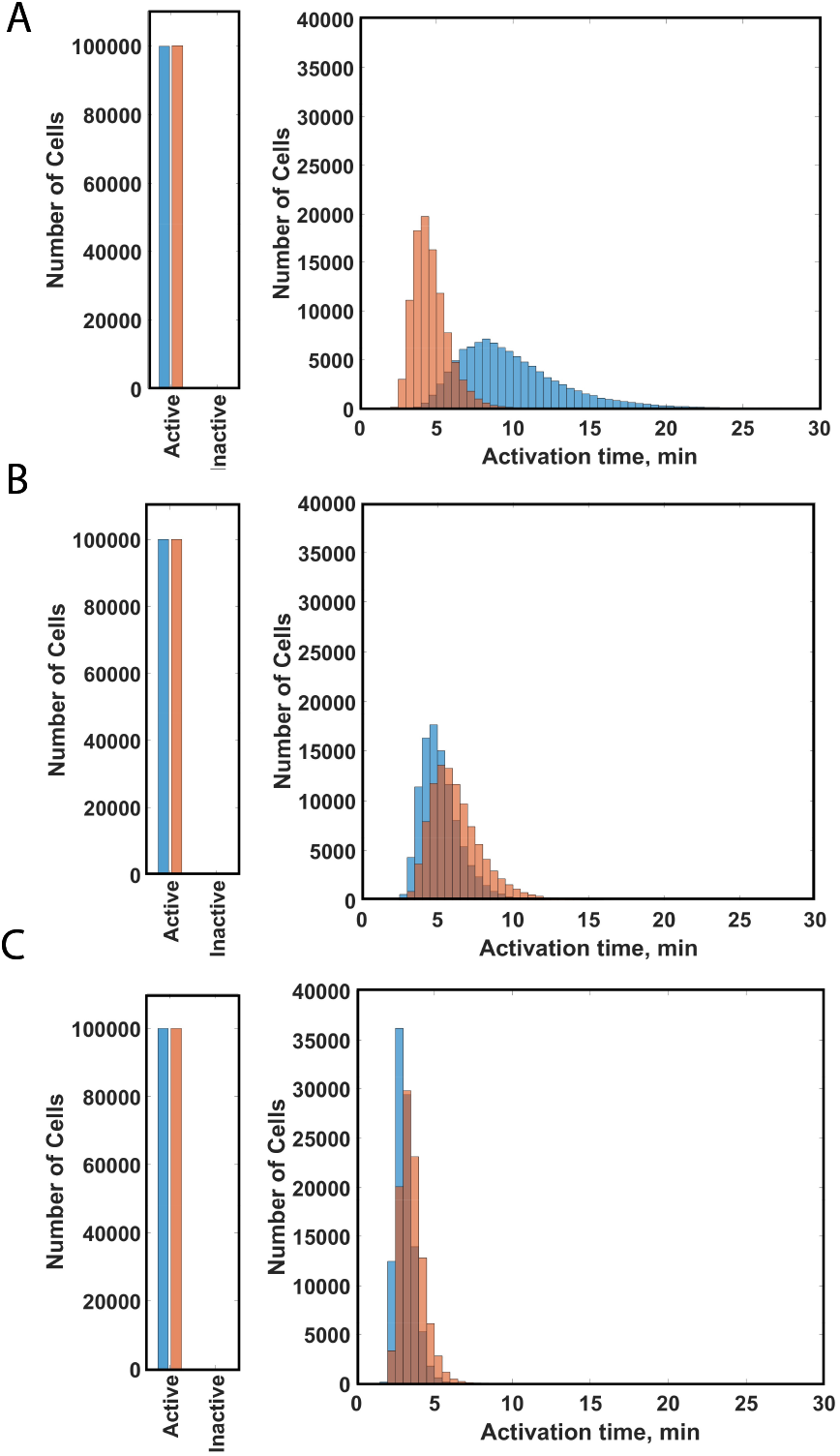
Activation of CAR T cells with optimized parameter values. We simulated simulation of cells with first- or second-generation CAR constructs upon implementing the optimized values of the two most influential parameters, *Kcat_ZAP* and *Kcat_LCKPU_CD3z*. (A) Simulated results with optimized *Kcat_ZAP* only; (B) Simulated results with optimized *Kcat_LCKPU_CD3z* only; (C) simulated results with both *Kcat_ZAP* and *Kcat_LCKPU_CD3z* optimized. *Blue*, predictions for cells with a first-generation CAR construct (CAR-CD3ζ); *orange*, predictions for cells with a second-generation CAR construct (CAR-CD3ζ-CD28).

**Figure 7.**
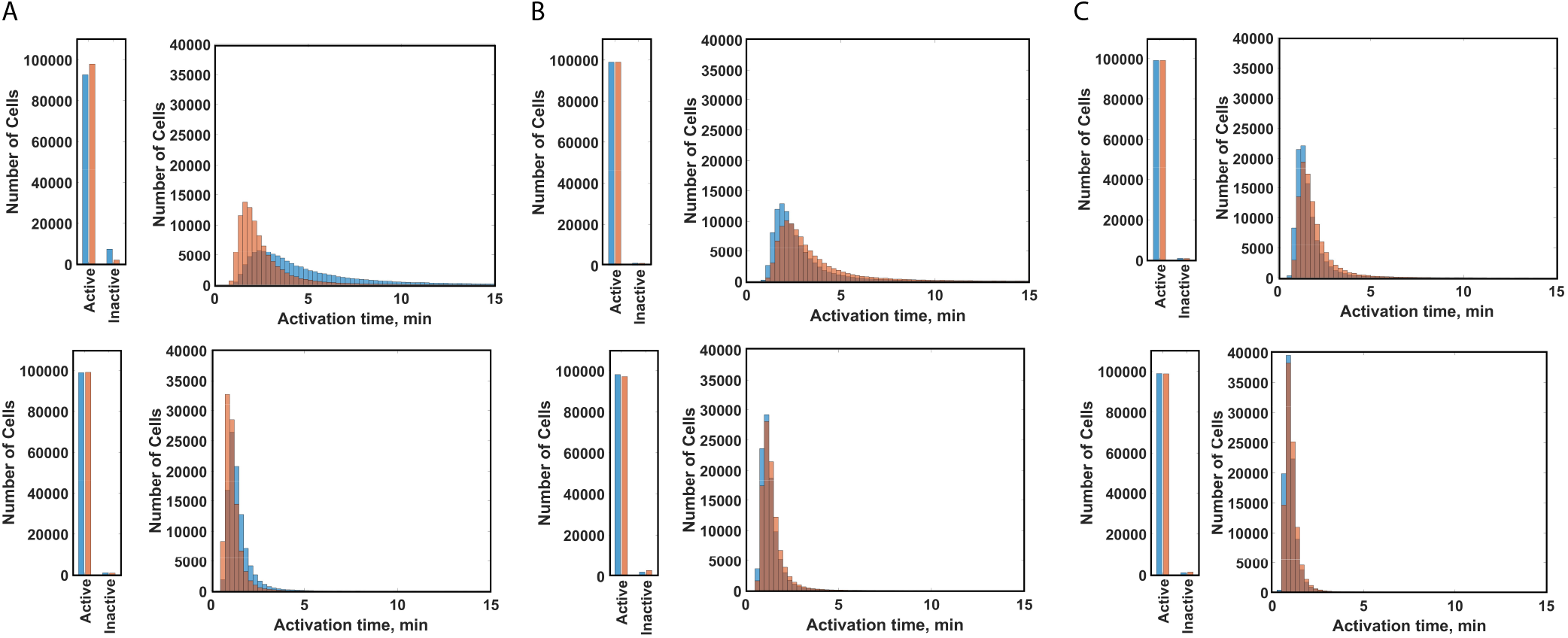
Activation of CAR T cells with varying kinetic parameters and optimized values of Kcat_ZAP and Kcat_LCKPU_CD3z. Simulated results are presented for low antigen concentration (top row) and high antigen concentration (bottom row). (A) *S*imulated results with optimized *Kcat_ZAP* only; (B) Simulated results with optimized *Kcat_LCKPU_CD3z* only; (C) Simulated results with both *Kcat_ZAP* and *Kcat_LCKPU_CD3z* optimized. *Blue*, predictions for cells with a first-generation CAR construct (CAR-CD3ζ); *orange*, predictions for cells with a second-generation CAR construct (CAR-CD3ζ-CD28).

## 4 DISCUSSION

In this study, we applied an existing ODE-based mechanistic model of CAR-induced ERK signaling to compare the performance of first- and second-generation engineered CAR-T cells given external and internal sources of variability. By simulating variable antigen concentrations, we showed that costimulatory signaling from CD28 causes both a reduced activation time and more consistent population response. In addition, by simulating stochastic kinetic interaction parameters, we were able to show that when considering kinetic variability caused by biochemical noise, second-generation cells yield similar results: reduced activation time and variability of the population response compared to first-generation cells. In order to track the mechanism by which CD28 induces this change, we repeated simulations of kinetic variability with only one CD28-mediated mechanism available at a time: the association between CD28 and Grb2, the association between CD28 and GADS, or the CD28-mediated change in the catalytic activity of LCK. We found that CD28’s effect on the catalytic activity of LCK is both necessary and sufficient to produce the performance improvement in second-generation cells. Next, we set out to identify parameters that could be modulated to further reduce response times and variability in the population response. To this end, we trained a gradient-boosted tree predictor, which is able to predict system activation time based on values of kinetic parameters. After confirming the predictor’s accuracy, we quantified the importance of each parameter in the predictor’s estimation of activation time. Using this method, we isolated five influential kinetic parameters. Then, we performed a constrained optimization procedure on our model by using an objective function that sought to minimize system activation time while manipulating as few of the five candidate parameters as possible. Based on results of this optimization procedure, the most optimal targets are predicted to be *Kcat_LCKPU_CD3z* and *Kcat_ZAP*. To elucidate how implementing the values suggested by the optimization procedure would affect population performance of the engineered T-cells, we repeated simulations of the system with antigen variability and kinetic variability by using the optimized parameter value. Optimizing *Kcat_ZAP* resulted in an overall reduction in cell response time but preserved the overall superiority of second-generation cells. In contrast, manipulating *Kcat_LCKPU_CD3z* independently or in conjunction with *Kcat_ZAP* not only drastically improved performance, but also made first-generation cells perform at least as efficiently as, and in some cases better than, second-generation cells.

Experimental studies have shown that the second-generation CAR constructs with a CD28 costimulatory domain promote a better immune response during *in vivo* testing, compared to first-generation constructs. While theoretical underpinnings of this phenomenon were explored in prior research, our work provides a new context for this discrepancy. Particularly, we showed that the incorporation of the CD28 costimulatory domain results in both shorter and more consistent response times in the face of various sources of variability. A population of CAR-T cells infused into the patient’s bloodstream would face highly variable external and internal conditions. As this variability is not reflected in the *in vitro* setting, our predictions provide valuable insight into the features of engineered CAR-T cells. In addition, we explored potential routes for the further improvement of CAR-T therapies. So far, work has mostly focused on incorporating more signaling domains or activity-dependent expression cassettes into the structure of the CAR (Akhoundi et al., 2021). We explored the alternative possibility of enhancing CAR T cell response by manipulating the catalytic activity of enzymes involved in signal transduction. The ability to engineer enzymes with desired properties, including improved catalytic activity, has already found broad applications in biotechnology (Lutz and Iamurri, 2018). While traditional methods use either a targeted substitution of key amino acids or directed evolution of random mutations, new approaches based on machine learning empower specialists to look for candidates *in silico* (Yang et al., 2019). Based on our model, engineering a more catalytically active isoform of LCK would cause first-generation cells to become at least as efficient as otherwise equivalent CD28-bearing CAR-T cells. It is established that despite their greater efficacy, second-generation cells suffer from multiple side effects, and mitigating those side effects is a significant concern. We believe that enhancing LCK catalytic activity as an alternative to having the CD28 domain in the CAR structure can be one such mitigating strategy, since it would potentially exclude undesirable effects of CD28 without compromising ERK signaling efficacy.

Along with the significant findings produced by our work, we recognize some limitations of our approach that can be addressed in the future. When exploring the population response of CAR-T cells to stochastic antigen concentrations, we assumed lognormal distribution parameters chosen to make the distribution fall within an experimentally observed range. For simulations of kinetic heterogeneity, we assumed a normal distribution centered around the accepted default value with a standard deviation as a third of this value. However, we have no unequivocal evidence that the assumed distributions in fact reflect what is observed experimentally. While our assumed distributions have a clear experimental basis both in terms of the distribution chosen and the parameters, they are not the only option possible. By trying other assumed distributions via altering the distribution parameters, a more comprehensive understanding of the CAR-T cell’s population response may emerge. Another limitation of our study concerns the choice of optimized values for *Kcat_ZAP* and *Kcat_LCKPU_CD3z* when considering heterogeneity. Our objective function penalized changes in each parameter proportional to the absolute value of the logarithm of the fold change compared to its baseline value. Since the baseline value of *Kcat_LCKPU_CD3z* is different between first- and second-generation cells due to CD28’s effects in the latter, we obtained different optimal values of *Kcat_LCKPU_CD3z* for first- and second-generation cells. However, since our ultimate goal was to simulate the observed effects of an artificially enhanced LCK, we assumed that the catalytic properties of such enhanced LCK would be the same regardless of the CAR structure. Thus, we were compelled to use the same optimal *Kcat_LCKPU_CD3z* value when performing simulations that accounted for antigen and/or kinetic variability. Future experimental work can explore the validity of our assumption.

## 5 CONCLUSIONS

Our work focused on exploring the difference in ERK activation times between engineered CAR-T cells with or without CD28 under various sources of cellular variability. We discovered that in line with expectations, the CD28 domain increases the proportion of cells that become activated based on ERK phosphorylation. Further, the model increases our mechanistic understanding of the role of CD28, predicting that CD28 confers shorter and more consistent activation times across the cell population. In addition, we discovered that the catalytic activity of LCK can serve as a valid target for the further improvement of CAR-T cell activation, since increasing the value of the corresponding parameter causes a pronounced improvement in the performance of first-generation CAR-T cells. Our work provides novel quantitative insights that can guide the design of CAR-engineered cells for immunotherapy.

## ACKNOWLEDGEMENTS

The authors thank members of the Finley research group for critical comments and suggestions.

## SUPPORTING INFORMATION

**File S1**. Supplementary figures and table (pdf file).

