## Supplementary Information for "Modeling predicts differences in CAR T cell signaling due to biological variability"

**Supplementary Table 1.** Performance of the gradient boosted tree ensemble on various datasets.

| Construct used: / Antigen conc.: | First-generation | Second-generation |
|---|---|---|
| "Low" | $R^2$ = 0.905±0.0008<br>EV = 0.905±0.0008 | $R^2$ = 0.900±0.0004<br>EV = 0.900±0.0004 |
| "High" | $R^2$ = 0.816±0.0024<br>EV = 0.816±0.0024 | $R^2$ = 0.807±0.0035<br>EV = 0.807±0.0035 |

### Supplementary Figures

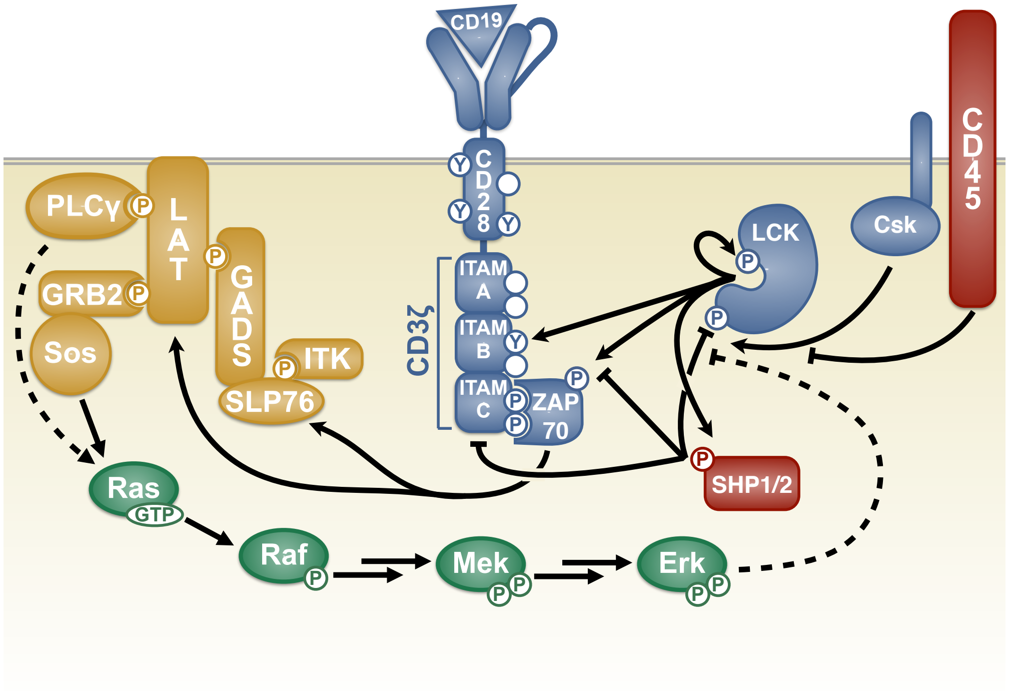


**Supplementary Figure 1. Schematic of CAR T cell signaling model.** We consider four modules. *Module I* (blue): CAR activation by LCK (whose catalytic activity is regulated by autophosphorylation and the inhibitory kinase, CSK). *Module II* (red): inhibition of CAR activation by phosphatases CD45 and SHP1/2. *Module III* (yellow): formation of the LAT signalosome, a multi-protein complex. *Module IV* (green): downstream signaling in the MAPK pathway, leading to ERK activation. Lines with bar indicate inhibition; lines with arrowheads indicate activation steps.

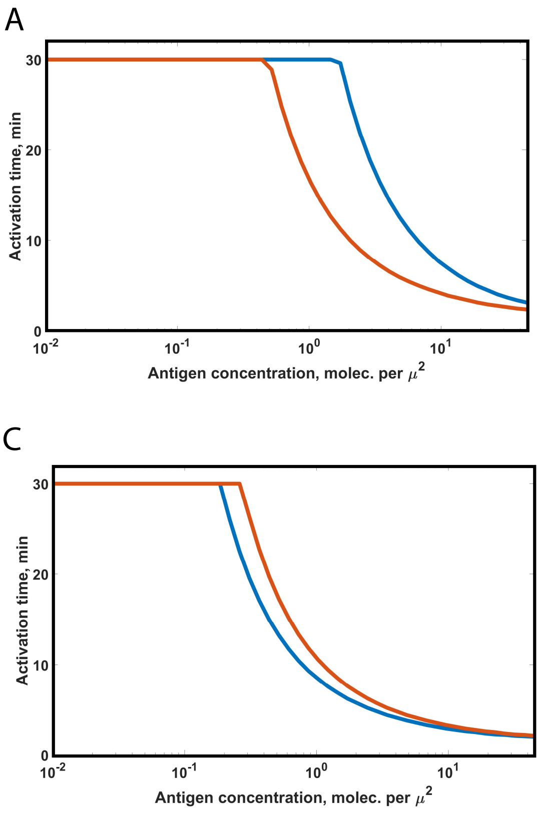


**Supplementary Figure 2. Dose response curves of cell activation times.** We varied the level of antigen exposure and simulated the activation time for different conditions. (A) Default parameters; (B) Optimized value for *Kcat_ZAP*, (C) Optimized value for *Kcat_LCKPU_CD3z*, (D) Optimized values for both *Kcat_ZAP* and *Kcat_LCKPU_CD3z*. *Blue*, predictions for cells with a first-generation CAR construct (CAR-CD3ζ); *orange*, predictions for cells with a second-generation CAR construct (CAR-CD3ζ-CD28).

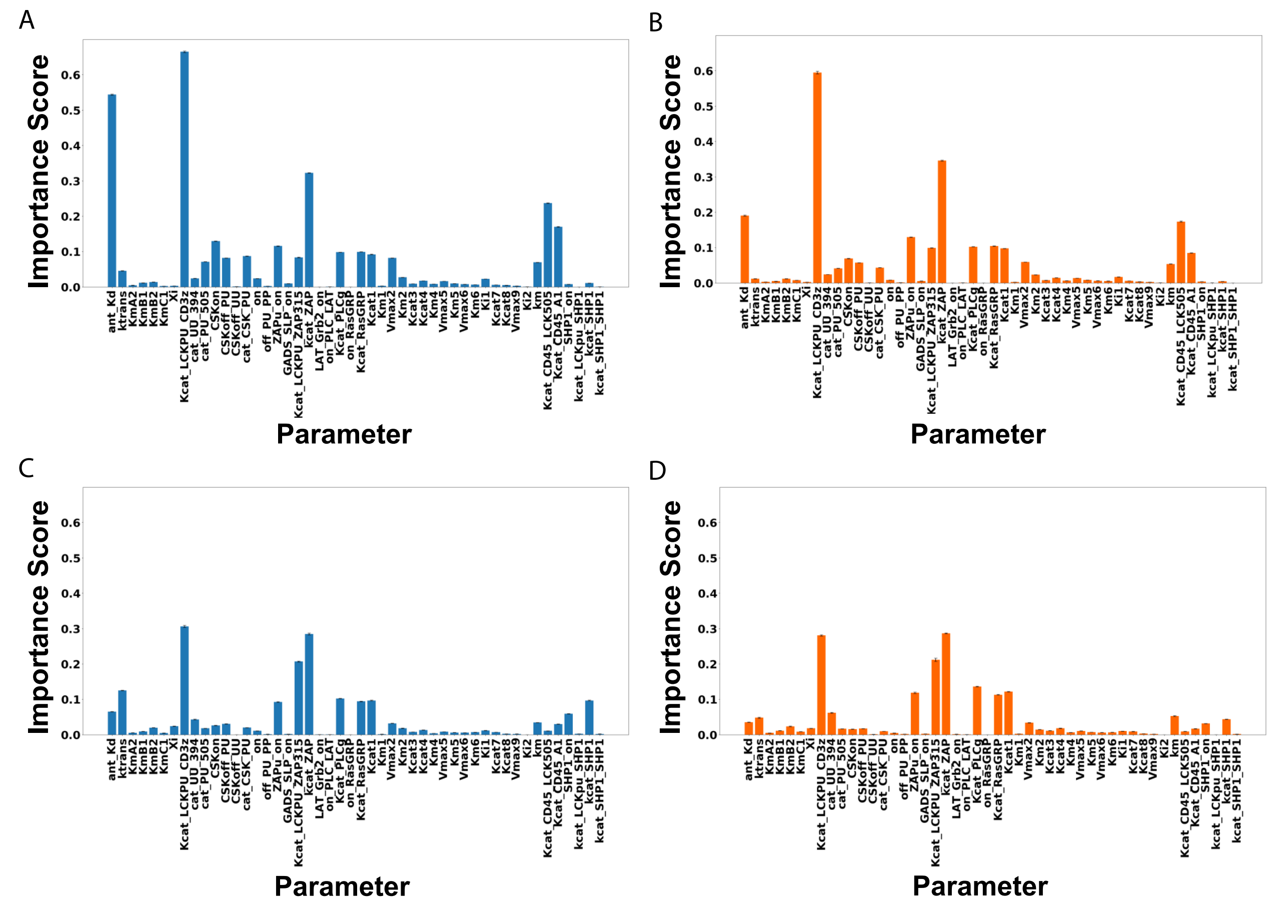


**Supplementary Figure 3. Permutation importance scores for 48 kinetic parameters used for creating a machine learning model to predict cell activation times.** A gradient boosted tree was used to predict the cell activation times based on model kinetic parameters. We show the permutation importance scores for all kinetic parameters under different conditions: (A) CAR-CD3ζ with low antigen concentration, (B) CAR-CD3ζ-CD28 with low antigen concentration, (C) CAR-CD3ζ with high antigen concentration, (D) CAR-CD3ζ-CD28 with high antigen concentration.

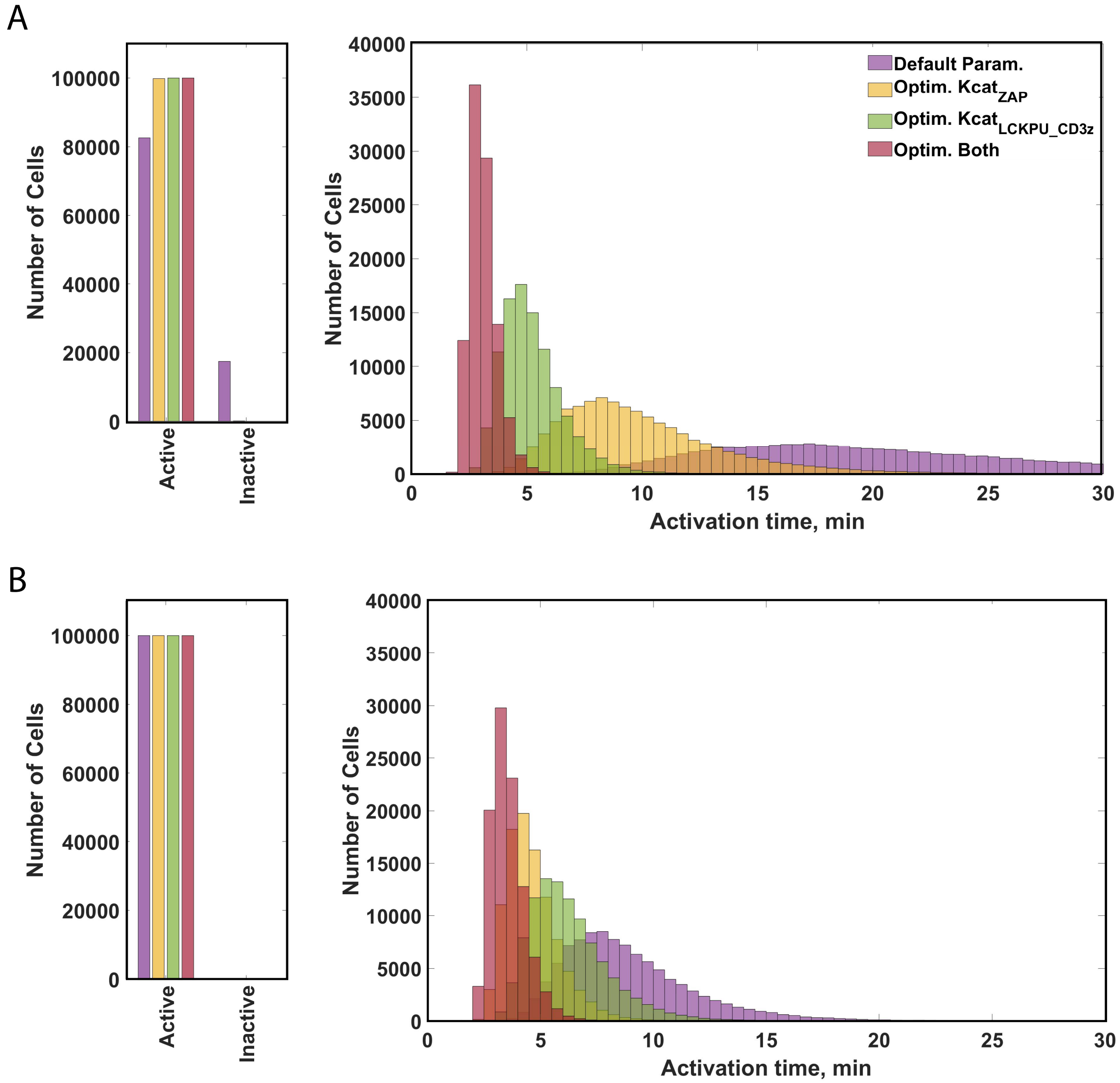


**Supplemental Figure 4. Comparison of activation of CAR T cells with optimized parameter values**. We show the number of cells activated and their activation times, showing shifts in the response of CAR-T cells with optimized kinetic parameters under antigen variation. Note, this is the same simulation data as shown in Fig. 1 and Fig. 6. (A) Comparison of the number of activated cells and their activation times for default and optimized parameter values among cells with first-generation CAR constructs (CAR-CD3ζ); (B) Comparison of the number of activated cells and their activation times for default and optimized parameter values among cells with second-generation CAR constructs (CAR-CD3ζ-CD28).

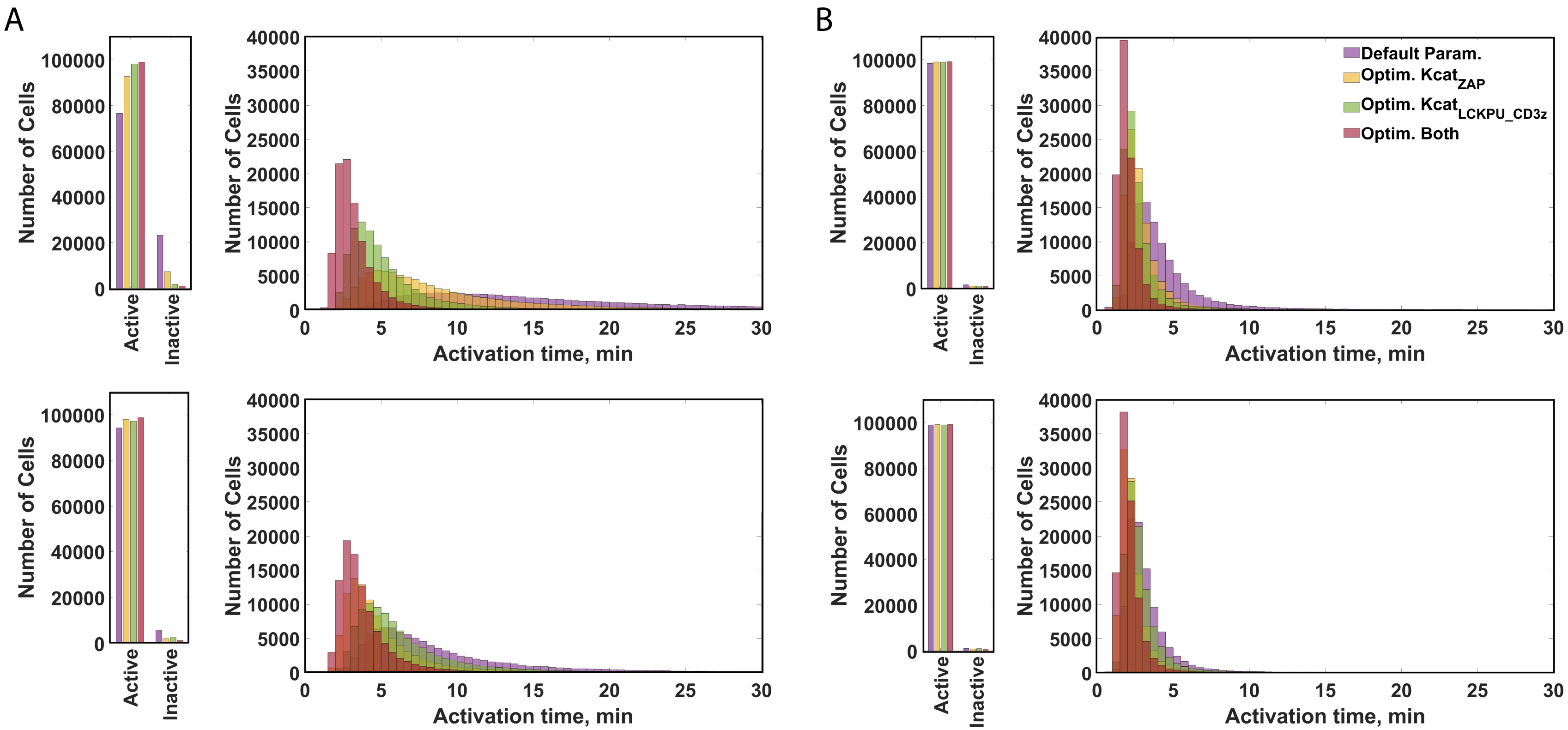


**Supplemental Figure 5. Comparison of activation of CAR T cells with optimized parameter values**. We show the number of cells activated and their activation times, showing shifts in the response of CAR-T cells with optimized kinetic parameters under kinetic parameter variation. Note, this is the same simulation data as shown in **Fig. 2 and Fig. 7**. (A) Comparison of the number of activated cells and their activation times for default and optimized parameter values among cells with first-generation CAR constructs (CAR-CD3ζ), top row: low antigen stimulation, bottom row: high antigen stimulation; (B) Comparison of the number of activated cells and their activation times for default and optimized parameter values among cells with second-generation CAR constructs (CAR-CD3ζ-CD28). *Top*: low antigen stimulation, *bottom*: high antigen stimulation.
